# Insights into the microevolution of SARS-ACE2 Interactions: In-silico analysis of glycosylation and SNP pattern

**DOI:** 10.1101/2022.06.29.498095

**Authors:** Pavan K Madasu, Arpita Maity, Surya K. Ghosh, Thyageshwar Chandran

## Abstract

The prefatory protein-glycan interaction and stabilizing protein-protein interaction of severe acute respiratory syndrome viruses with angiotensin-converting enzyme 2 play a significant role in complex formation thereby promoting endocytosis. The microevolution of SARS-CoV-2 over a period of time has a significant role in increasing the affinity of receptor-binding domain against angiotensin converting-enzyme 2. In the current study, we have corroborated the vitality of acquired SNPs over a period of time with increased affinity by using docking studies. The results indicate that the virus modulates the undesirable glycosylation sites by a series of substitution and deletion mutations. It uses bulky residues such as Tyr/Phe for dynamic arrest for quick stabilization of the complex, and Lys residues for stabilizing via hydrogen bond formation besides increasing the binding affinity to ease the cell entry.

## Introduction

The spike protein of the severe acute respiratory syndrome – Corona virus-2 (SARS-CoV-2) exhibits two types of interactions in infecting its host cells. One is the usual protein-protein interactions (PPI) and the other is the protein-glycan interactions (PGI). The PPI ensures the formation of a stable complex of spike protein with angiotensin converting enzyme 2 (ACE2) by electrostatic forces, hydrogen bonding, and hydrophobic effect (Agamennone et al., 2021; Jones & Thornton, 1995) whereas PGI serves as the preliminary point of contact for priming the interactions (Gimeno et al., 2020; Raman et al., 2016). In the case of the recent variant of SARS-CoV-2 (Omicron), the 452-498 loop in the receptor-binding domain (RBD) of spike protein plays a significant role in PPI(Baral et al., 2021). The mutations (Q495R and N498Y) in the said loop were associated with increased binding affinity and the same was confirmed by *in-vitro* evolution studies (Zahradník et al., 2021).. Recent studies indicate the role of two more SNPs (S477N:Omicron ; T478K:Omicron and Delta) in functional gain mutations. In the case of S→N, the virus gain an additional N-glycosylation site, while the later led to increased binding affinity by means of hydrophobic interactions (Cherian et al., 2021; Singh et al., 2021). Intriguingly, the D614G substitution in the S1/S2 cleavage site which was first reported in Beta variant has been carry forwarded to the subsequent variants increasing flexibility of the S2 subunit thereby improving the ability of viral membrane protein to fuse with the cell membrane (Cherian et al., 2021; Ozono et al., 2021). In the current study, we have corroborated the increased stability of the RBD-ACE2 complex formation favored by PPIs of the acquired SNPs by docking studies of notable SARS variants with ACE2.

The PGI of the SARS with ACE2 is established by the glycans distributed over the protein surface. The liquid chromatography-mass spectroscopic (LC-MS) studies of glycosylated spike protein confirm the presence of 22 N-glycosylation and 2 O-glycosylation sites (Shajahan, Supekar, et al., 2020; Watanabe et al., 2020). Research evidence suggests that N-glycans N165 and N234 play an essential role in modulating the conformational dynamics of the receptor-binding domain (RBD) thereby influencing ACE2 recognition (Casalino et al., 2020) whereas reduced infectivity is associated with deletion of N331 and N343 glycosylation sites (Li et al., 2020). In the current study, we have emphasized the evolutionary pattern of N-and O-glycan sites for PGI interactions among the SARS variants.

## Materials and Methods

### Data retrieval

The SARS-CoV-2 spike glycoprotein sequence data was retrieved from NCBI updated by Phylogenetic Assignment of Named Global Outbreak LINeages (pangolin) web app maintained by Centre for Genomic Pathogen and Surveillance. The SARS-CoV-2 variants and mutations data was retrieved from the data (https://covariants.org/) curated by the Institute of Social and Preventive Medicine University of Bern, Switzerland and SIB Swiss Institute of Bioinformatics, Switzerland. The new double mutant variant with all the Omicron SNPs clubbed with deletion loop (246-252) of Lambda and Delta are manually curated and named as Lamicron and Delmicron variant.

### Single Nucleotide Polymorphisms analysis

The SARS-CoV-2 variants and mutations data of 22 variants (https://covariants.org/) were scrutinized for the recurring ones among all the variants. Further, the nature of the variants was analyzed by using the consensus tool PredictSNP (Bendl et al., 2014). The SNPs from all the variants are listed and ranked according to their periodicity. The percentage of the periodicity was calculated by the formula:

Percentage = SNP found in number of variants / Total number of variants X 100

### Molecular Evolutionary and Phylogenetic analysis

The 3D spike glycoprotein structures of all the SARS variants were developed using Modeller 9.23 (Webb & Sali, 2016). The multiple sequence alignment of all the SARS variants including SARS-CoV was carried out using structure based sequence alignment TCOFFEE (Armougom et al., 2006). The microevolutionary pattern was inferred using the minimum evolution method in MEGA X (Kumar et al., 2018) using Close-Neighbor-Interchange (CNI) algorithm.

### Prediction of Glycosylation Sites

The N-and O-linked probable glycosylation sites of all the SARS variants were envisaged by using GLYCAM glycoprotein builder (http://legacy.glycam.org/tools/molecular-dynamics/glycoprotein-builder/upload-pdb) considering the threshold solvent accessible surface area (SASA) as 40.

### Molecular Docking and Analysis

The flexible protein-protein docking of selected SARS variants (SARS-CoV, SARS-CoV-2, Lambda, Omicron, Lamicron and Delmicron) with angiotensin-converting enzyme 2 (ACE2) was performed using High Ambiguity Driven protein-protein DOCKing (HADDOCK) server (Honorato et al., 2021). The binding affinity (Δ G=-RTlnK_d_) at 25^0^C of the RBD-ACE2 complex generated by HADDOCK were analyzed using the PRODIGY server (Xue et al., 2016).

The angle of orientation of RBD of the SARS variants with ACE2 receptor of *Homo sapiens* was calculated by considering centre of mass of ACE2, RBD with 325-329 loop of ACE2 as midpoint.

## Results and Discussion

The spike glycoprotein sequence data of 9 variants (Omicron, Eta, Iota, Kappa, Lambda, Gamma, 20E EU1, 20B/S, and Delta) were successfully retrieved from NCBI database. Additionally, based on the mutation data available on CoVariants webpage (https://covariants.org/) a total of 13 SARS-CoV-2 variants (Alpha, Beta, Mu, 20 A/S 126A, 21C Epsilon, 20 A/S 439K, S677H Robin1, S677P Pelican, 20A EU2, 20A/S 98F, 20C/S 80Y, 20B/S 626S, and 20B/S 1122L) were manually generated, taking the total available variants to 22. A total of 93 SNPs could be observed from the said variants, of which 13 were found to be the most recurring ones. D614G SNP was found in 15 out of 22 variants with a periodicity of 68.2 % ; E484A/K/Q SNP has periodicity 8/22 (36.36%); P681H/R SNP has periodicity 7/22 (31.81%); N501Y SNPs have periodicity 5/22 (22.72%); H69-, V70-and Y144-/S have periodicity 4/22 (18.18%); L452R/Q, K417N/T, S477N, T478K, T95I and H655Y has periodicity 3/22 (13.63%) whereas all other 81 SNPs have periodicity ≤ 2/22, respectively(Table S1). Out of 93 SNP substitutions, 12 (Q52R, T95I, W152C, R190S, L452Q/R, Y505H, N764K, N856K, F888L, N969K, T1027I, and D1118H) were classified as deleterious based on the PredictSNP prediction (Bendl et al., 2014) (Table S2) and all these SNPs have a periodicity ≤ 3/23.

The structure based sequence alignment of all the SARS variants showed 99% similarity (data not shown). The active site residues (Table1) of all the variants are tabulated considering data from the decade long structural analysis of SARS Variants from various hosts (Wan et al., 2020). The microevolution of SARS variants over a period of time confirms that the Omicron variant completely differs from all other variants evolved in a short period of time (Figure 1). Intriguingly, compared with other variants it possess a high number of SNPs (35) of which all the functionally significant SNPs were found to be conserved. The most recent variant Omicron 21K also has 35 SNP sites (Figure 2), of which 2 (6%) have acquired the N-glycan site, 2 (6%) the O-glycan site and 5 (14%) bulky residues. The other Omicron 21L variant shares 22 common SNP sites with above mentioned variant. Additional N-glycan (D405N), O-glycan (A27S) and bulky residue (S371F) could also be observed.

**Figure 1:**
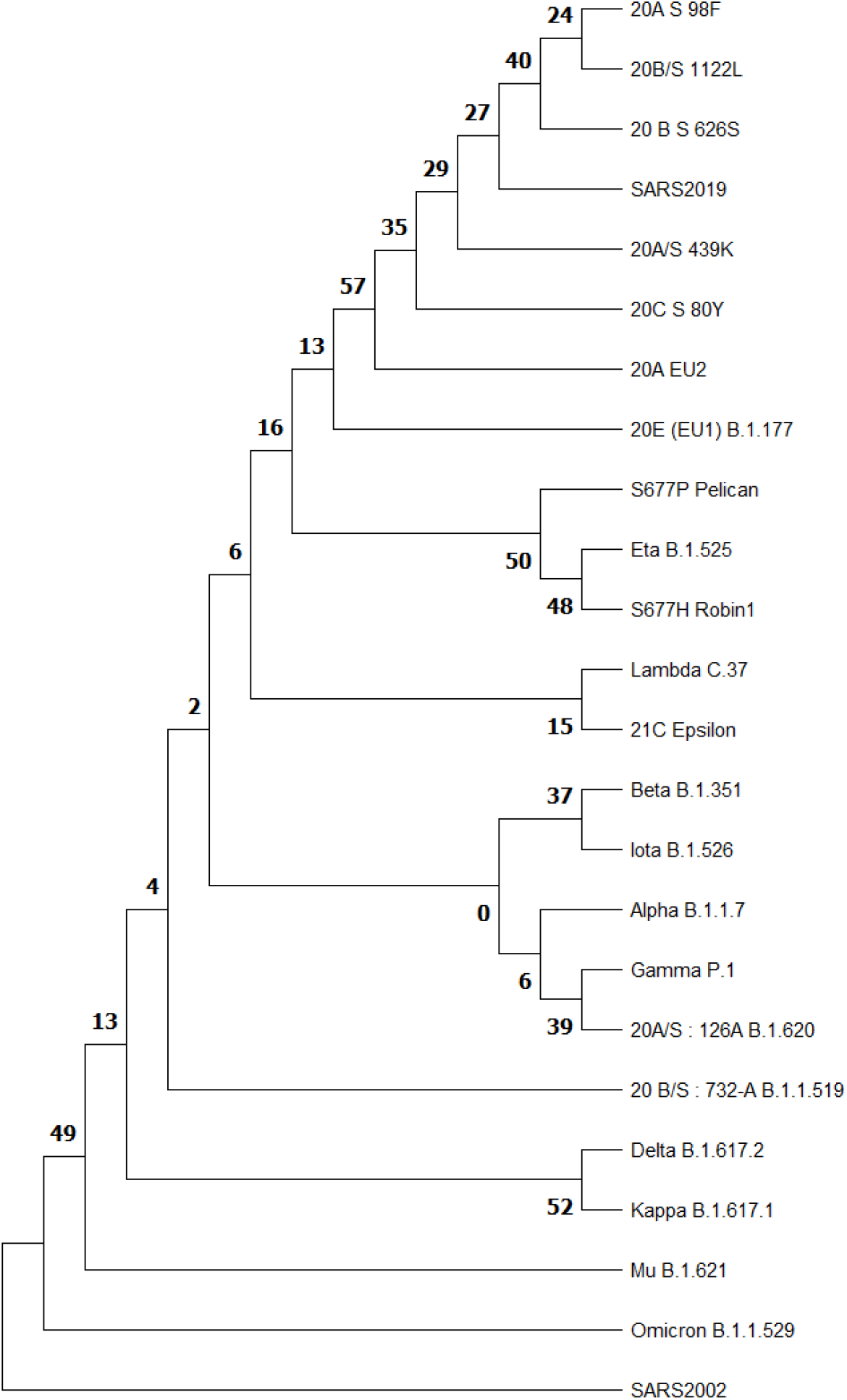
SNP Phylogeny of SARS Variants depicting the micro-evolution. (The percentage of replicate trees in which the associated taxa clustered together in the bootstrap test (1000 replicates) are shown next to the branches)

**Figure 2:**
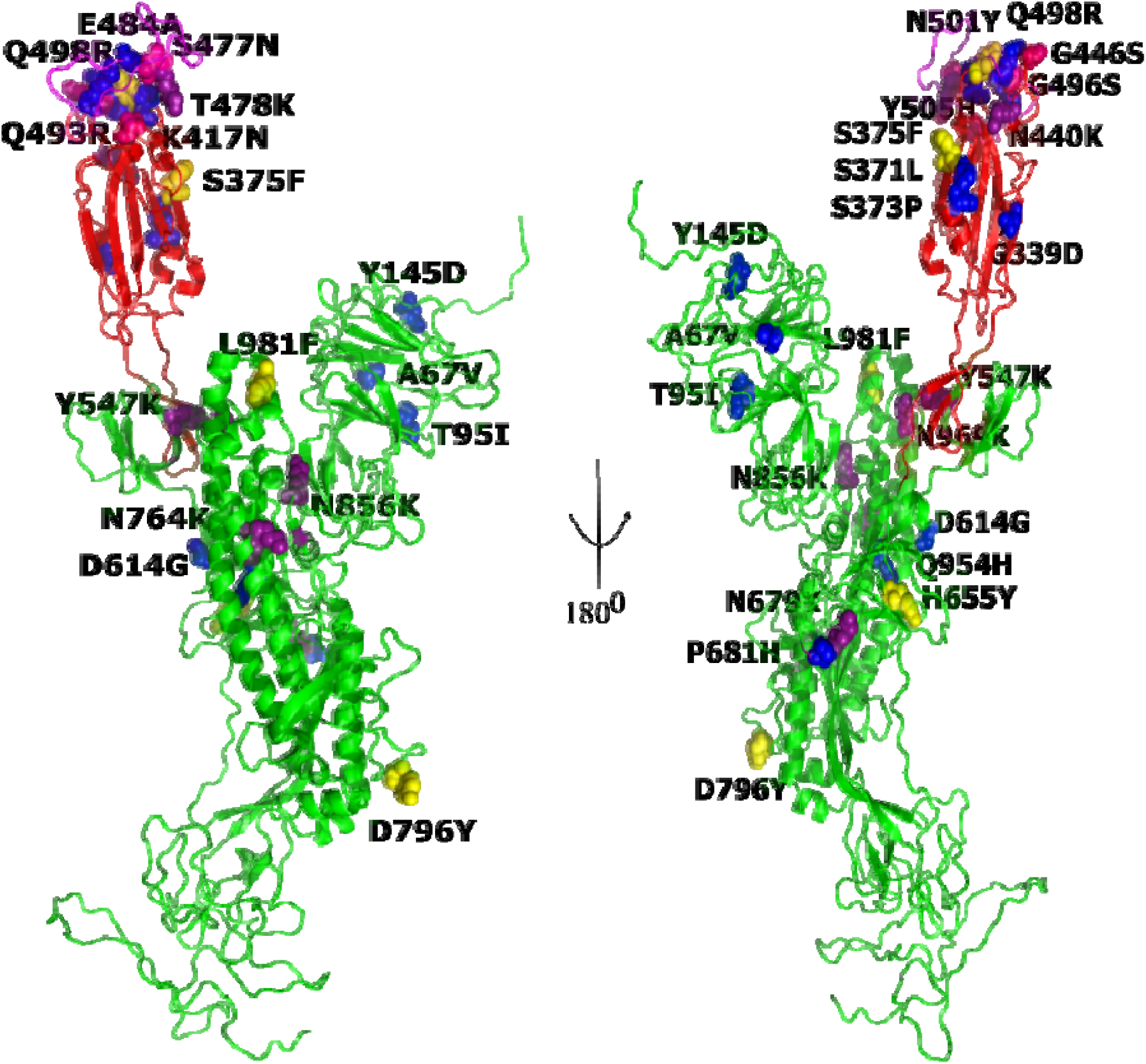
Single nucleotide polymorphism (SNP) sites of Omicron variant are shown in ball representation. The mutants are differently coloured (Aromatic-Yellow, Lys-deep purple, Glycosylation sites-magenta, and rest all-blue)

The number of N-and O-glycosylation sites observed in the complete spike glycoprotein and receptor-binding domain (RBD) are shown in Figure 3. There is an addition of 32 (16 N-linked & 16 O-linked) novel glycosylation sites from SARS-CoV →SARS-CoV-2 variant and thereafter significant deletion of a few sites were observed among the later variants. This deletion was found to be associated with the introduction of Lys (AAA/AAG) which is required for stabilizing via hydrogen bond formation as it is just one mutational space away from both Asn (AAT/AAC) and Thr (ACA/ACG). Moreover, the most prominent SNP to be considered in the evolutionary analysis is T501N (SARS-CoV→SARS-CoV-2) in the interacting loop of RBD i.e., change in O-glycosylation site to N-glycosylation site. The N-glycan residue (Asn:AAT/AAC) is just one nucleotide distance away from both the O-glycan residues (Ser:AGT/AGC and Thr:ACT;ACC) which indicates the preference of N over O glycosylation. Recent LC-MS glycosylation profile studies (Shajahan et al., 2020; Watanabe et al., 2020) have confirmed the same. Later on, this site mutated to Tyr creating an N501Y variant; the probable reason for this mutation could be attributed to the very low SASA (17.5) score for Asn which cannot accommodate any glycans on the said site. Eventually, under evolutionary pressure the said site was mutated to a bulky aromatic residue, providing the necessary stacking interactions besides arresting the dynamics of RBD, inturn increasing the binding efficiency of RBD-ACE2. The probable justification for this substitution is that the Asn (AAT/AAC) is just one nucleotide away from Tyr (TAT/TAC). Most of the SNPs acquired by the recent SARS variants have Lys, Arg, and Asp substitutions which are known to provide stacking and stabilizing interactions via hydrogen bonds. The mutational probability of Asp to Tyr/Gly/Arg and Arg to Gln/Lys/Pro is high as they are just one nucleotide distance away (Table S1). In order to understand the effect of SNPs on the binding dynamics of SARS variants RBD domain with ACE2, we carried out modelling studies as and when required.

**Figure 3:**
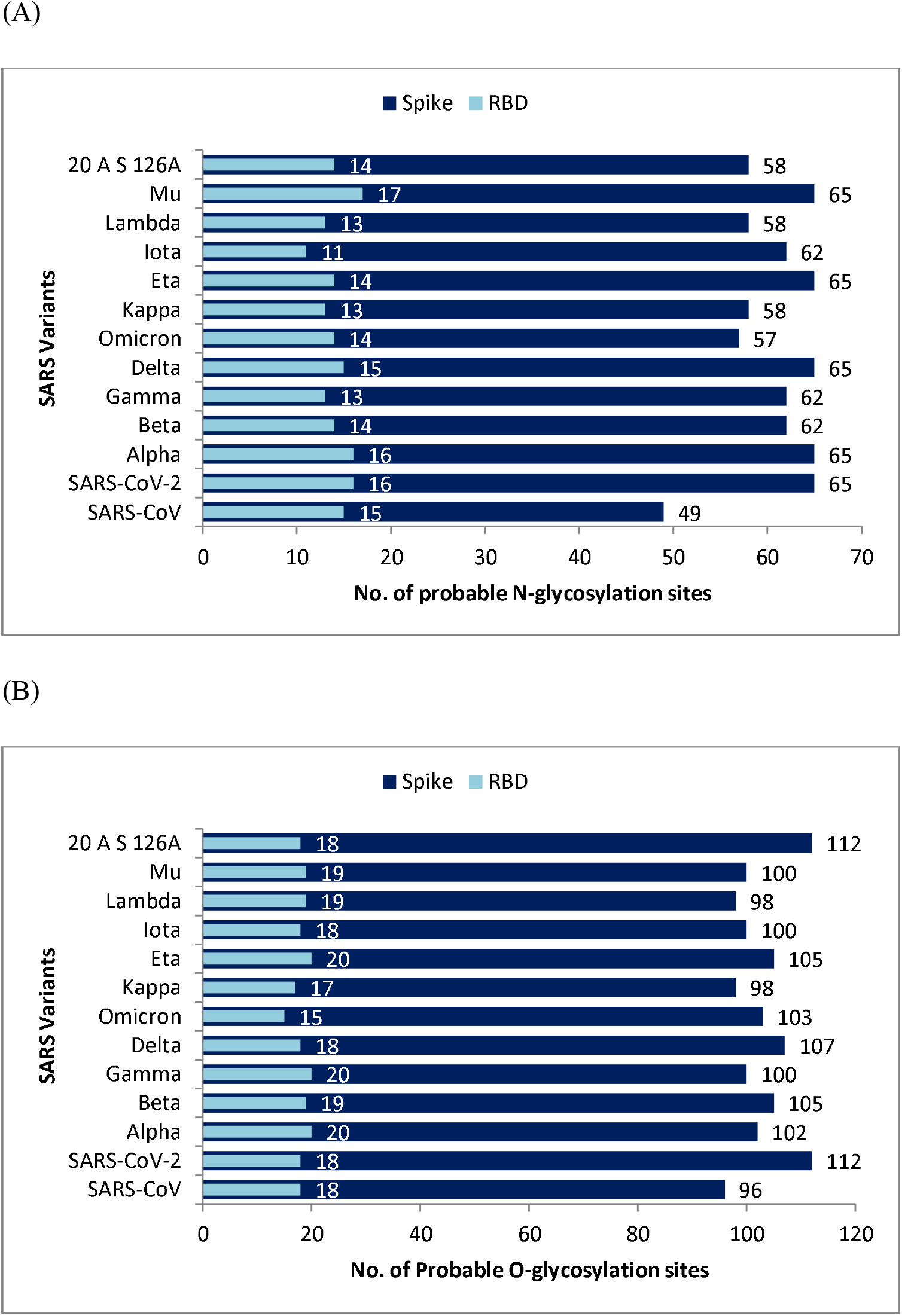
Predicted (A) N-glycosylation and (B) O-glycosylation sites in Spike protein (dark blue) and RBD (sky blue) in bar in bar representation. The X-axis represents the No. of probable sites and the Y-axis represents the SARS Variants arranged from bottom according to their occurrence.

The modelled S-protein from different variants were then docked onto ACE-2 receptor. The docking studies (Figure 4) confirm that the affinity of S protein towards ACE2 has increased over time for the variants with one exception of SARS-CoV-2 (Table 2). Intriguingly, for the SARS-CoV-2 (Δ G = -9.5 Kcal/mol & Kd = 1.0E-07 M) variant has decreased affinity compared with its precursor SARS-CoV (Δ G = -9.9 Kcal/mol) variant which has a dissociation constant (Kd) of 5.8E-08 M. The prominent later variants Delta (ΔG = -10.5 Kcal/mol & Kd = 1.8E-08M), Lambda (ΔG = -9.8 Kcal/mol & Kd = 6.2E-08 M) and Omicron (ΔG = -11.4 Kcal/mol & Kd = 4.4E-09 M) have shown increased binding affinity towards ACE2 receptor respectively. Interestingly, the manually curated double mutant Lamicron (Lambda + Omicron) has shown decreased affinity (ΔG = -8.5 Kcal/mol & Kd = 5.9E-07 M) and can be correlated with non-conservation of the deletion loop (246-252) in later variants. On contrary, the double mutant Delmicron (Delta + Omicron) has shown increased binding affinity (ΔG = -12.2 Kcal/mol) while the dissociation constant was Kd = 1.2E-09 M. This indicates, if there is any new variant identical to delmicron then it will be more catastrophic than the existing variants.

**Figure 4:**
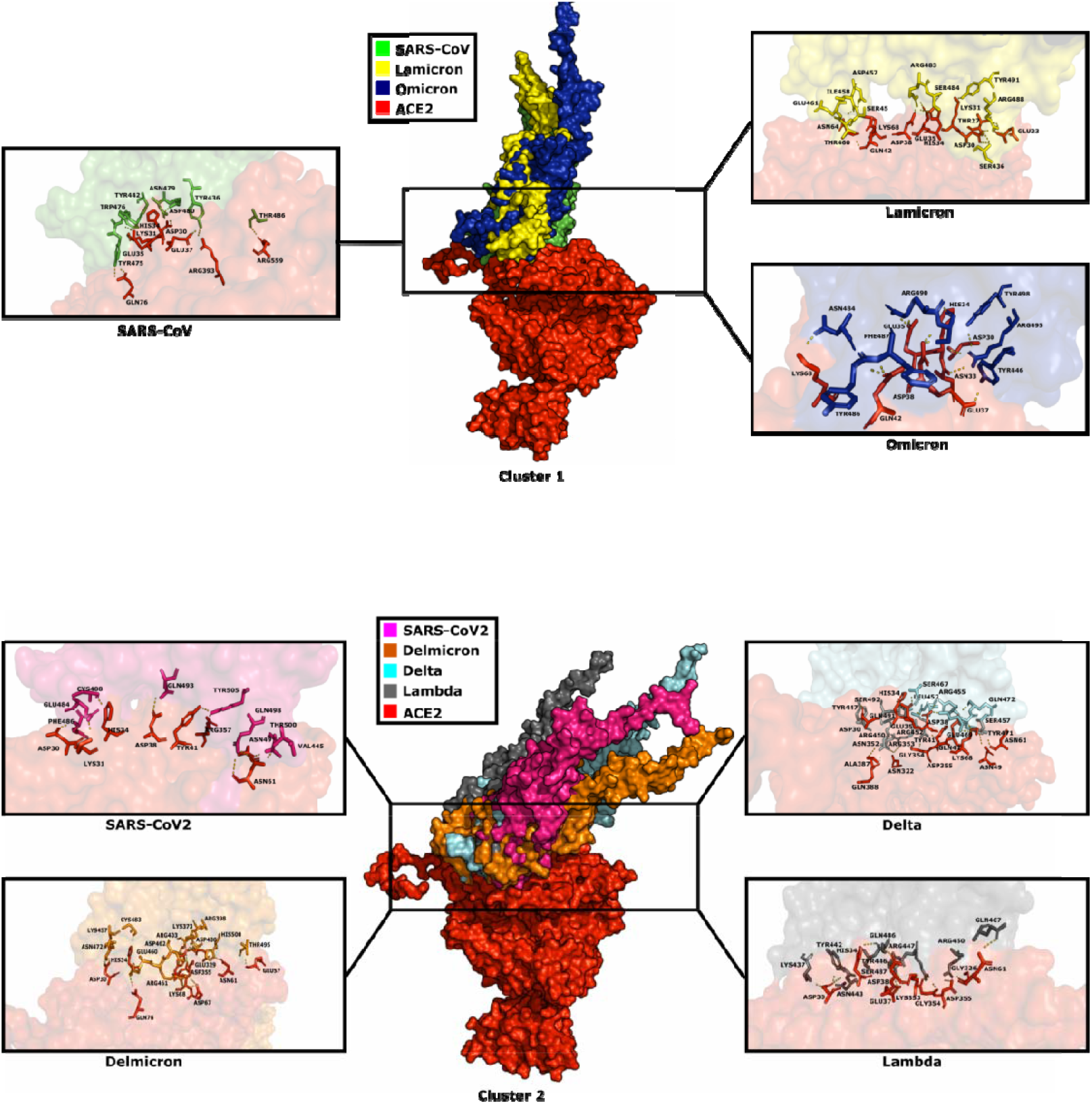
Grouping of SARS Variants into Cluster 1 and Cluster 2 based on their docking orientation with ACE2 receptor (red colour) of *Homo sapiens*. (The RBD of SARS variants are shown in surface representation and visualized in different colours: SARS-CoV (green); Lamicron (yellow); Omicron (blue); SARS-CoV-2 (pink); Delmicron (orange); Delta (Cyan) and Lambda (gray).

**Table 1:** Active site residues (numbers) found in the interacting loop of the reported SARS Variants. (including manually curated double mutants Lamicron and Delmicron)

| Variant | L | F | Q | S | Y |
| --- | --- | --- | --- | --- | --- |
| SARS-CoV | Y (442) | L (472) | N (479) | D (480) | T (487) |
| SARS-CoV-2 | 455 | 486 | 493 | 494 | N(501) |
| Alpha | 452 | 487 | 490 | 491 | 498 |
| Beta | 452 | 487 | 490 | 491 | 498 |
| Gamma | 455 | 490 | 493 | 494 | 501 |
| Delta | 453 | 488 | 491 | 492 | N (499) |
| Omicron | 452 | 484 | 490 | 491 | 498 |
| Lambda | 448 | S (483) | 486 | 487 | N (494) |
| Mu | 456 | 491 | 494 | 495 | 506 |
| Kappa | 455 | 490 | 493 | 494 | N (501) |
| Eta | 452 | 487 | 490 | 491 | N (498) |
| Iota | 455 | 490 | 493 | 494 | N (501) |
| 20B/S 732A | 455 | 490 | 493 | 494 | N (501) |
| 20A/S 126A | 449 | 484 | 487 | 488 | N (495) |
| 20E EU1 | 455 | 490 | 493 | 494 | N (501) |
| 21C Epsilon | 455 | 490 | 493 | 494 | N (501) |
| 20A/S 439K | 453 | 488 | 491 | 492 | N (499) |
| S677H Robin1 | 455 | 490 | 493 | 494 | N (501) |
| S677P Pelican | 455 | 490 | 493 | 494 | N (501) |
| 20A EU2 | 455 | 490 | 493 | 494 | N (501) |
| 20A/S 98F | 455 | 490 | 493 | 494 | N (501) |
| 20C/S 80Y | 455 | 490 | 493 | 494 | N (501) |
| 20B/S 626S | 455 | 490 | 493 | 494 | N (501) |
| 20B/S 1122L | 455 | 490 | 493 | 494 | N (501) |
| Delmicron | 450 | 485 | R (488) | 489 | 496 |
| Lamicron | 445 | 480 | R (483) | 484 | 491 |

**Table 2:** Binding affinity (ΔG) and disassociation constant (K_d_) at 25^0^C of RBD-ACE2 interactions of selected SARS Variants calculated using PRODIGY tool.

| **Interacting Protein Complex** | **$\Delta G$**<br>**(Kcal/mol)** | **$K_d$ (M) at 25<sup>0</sup>C** |
| --- | --- | --- |
| SARS CoV RBD-ACE2 | −9.9 | 5.8E-08 |
| SARS CoV 2 RBD-ACE2 | −9.5 | 1.0E-07 |
| Delta RBD-ACE2 | −10.5 | 1.8E-08 |
| Lambda RBD-ACE2 | −9.8 | 6.2E-08 |
| Omicron RBD-ACE2 | −11.4 | 4.4E-09 |
| Delmicron RBD-ACE2 | −12.2 | 1.2E-09 |
| Lamicron RBD – ACE2 | −8.5 | 5.9E-07 |

**Table 3:** Comparative set of residues forming H-bond table of notable SARS Variants with ACE2 receptor of *Homo sapiens*.

| ACE2 Receptor | SARS Spike Protein Variants |  |  |  |  |  |  |
| --- | --- | --- | --- | --- | --- | --- | --- |
|  | SARS CoV | SARS CoV 2 | Delta | Lambda | Omicron | Delmicron | Lamicron |
| Glu23 |  |  |  |  |  |  | Arg488 |
| Thr27 |  |  |  |  |  |  | Arg488 |
| Asp30 |  | Phe486 | Tyr447 | Lys437<br>Tyr442 | Arg495<br>Tyr498 | Lys457<br>Asn472 | Ser436<br>Arg488 |
| Lys31 | Tyr442 | Glu484 |  |  |  | Glu460 | Tyr491 |
| Asn33 |  |  |  |  | Arg495 |  |  |
| His34 | Asp480 | Cys488 | Gln491<br>Ser492 | Gln486 | Ser491 | Cys483 | Ser484 |
| Glu35 | Trp476 |  | Ser467 |  | Arg490 |  | Arg483 |
| Glu37 | Tyr436 |  |  | Tyr446 | Tyr446 |  |  |
| Asp38 |  | Gln493 | Gln491<br>Arg455 | Gln486 | Ser491 |  | Arg483 |
| Tyr41 |  | Tyr505 | Arg455 |  |  | Asp400<br>Arg403 |  |
| Gln42 |  |  | Gln472 |  | Tyr486<br>Phe487 |  | Thr460 |
| Asn49 |  | Thr500<br>Gln498 | Ser457 |  |  |  |  |
| Glu57 |  |  |  |  |  | Thr495 |  |
| Asn61 |  | Val445<br>Gln498 | Tyr471 | Gln467 |  | His500 |  |
| Asn64 |  |  |  |  |  |  | Ile458<br>Ser459 |
| Asp67 |  |  |  |  |  | Arg461 |  |
| Lys68 |  |  | Glu469 |  | Asn484 | Asp462 | Asp457 |
| Gln76 | Tyr475 |  |  |  |  | Arg461 |  |
| Asn322 |  |  | Asn352<br>Arg353 |  |  |  |  |
| Gly326 |  |  |  | Arg450 |  |  |  |
| Glu329 |  |  |  |  |  | Lys373<br>Arg398 |  |
| Lys353 |  |  |  | Arg447 |  |  |  |
| Gly354 |  |  | Tyr41 | Arg447 |  |  |  |
| Asp355 |  |  | Tyr41 | Arg450 |  | Arg403 |  |
| Ala387 |  |  | Asn352 |  |  |  |  |
| Gln388 |  |  | Asn352 |  |  |  |  |
| Arg393 | Tyr436 |  |  |  |  |  |  |
| Arg559 | Thr486 |  |  |  |  |  |  |

In order to further understand the binding mechanism between the RBD of the SARS Variants with ACE2 receptor of *Homo sapiens*, the angle of orientation was calculated (Table S3) which was found to be significantly decreased from SARS-CoV (114.9^0^) → SARS-CoV-2 (91.3^0^) with slight augment in the values for the later variants. This indicates that the virus is trying to evolve by a series of mutations to find the best docking angle with ACE2. We have clustered the above variants into two groups based on their docking orientation with ACE2 (Figure 4). The cluster 1 includes SARS-CoV, Omicron and Lamicron whereas the cluster 2 includes SARS-CoV-2, Delta, Lambda and Delmicron. The RBD binding site of all the SARS variants with ACE2 falls in the same region (Figure 4). The RBD domain is in the region of 306-540 aa where the loop 436-507 aa plays vital role in binding with ACE2 receptor (Table 5). A series of tinkering mutations could be observed within this loop. On the whole, 13 mutations (Table S1) were observed in this region, of which 5 (E484A/K/Q, N501Y, L452R/Q, S477N, T478K) of them have periodicity ≥ 3. The variants with the SNP N501Y are extremely transmissible and were considered as Variants of Concern (VOC) by Centre for Disease Control and Prevention (CDC) (https://www.cdc.gov/coronavirus/2019-ncov/variants/variant-classifications.html). Intriguingly, the SNP E484A/K/Q is the most recurring mutation among all the variants after D614G. Further replacement of this site with Arg(R) or Asp (D) may increase the binding affinity and is expected to be observed in the new upcoming variants (Cherian et al., 2021; Yang et al., 2022). The other 3 SNPs (L452R/Q, S477N, T478K) introduce additional Nitrogen atom required for the formation of hydrogen bonds with ACE2 leading to stabilized interactions. Additionally, 8 SNPs (S438K, N440K, G446S, F490S, Q493R, G496S, Q498R and Y505H) contribute the Nitrogen/Oxygen atoms required for the hydrogen bond formation. Further in-vitro/in-vivo dynamic studies will help us to understand whether the said SNPs augment the RBD-ACE2 interactions.

## Conclusion

During the course of evolution, SARS variants have gained a maximum number of N-and O-glycosylation sites (16 N and 16 O) but interestingly, it seems that the virus is getting rid of non-specific N-and O-glycosylation sites (Figure 2) in order to escape immune pressure from host and increase its virulence. As it happened in the case of Omicron, the substitution of Asn (N-glycosylation site) with bulky Tyr residue in the interacting loop led to the evolution of the variant of concern (VOC) N501Y variant. This confirms that glycosylation sites with less SASA are replaced with either bulky Tyr (N501Y of Omicron variant) residues for stacking interactions and Lys (N440K, T478K, T547K, N679K, N764K, N856K, and N969K) for increased hydrogen bond interactions. A potential glycosylation site should have a SASA of 40 less than which the site cannot accommodate glycans likely due to steric hindrances. The detailed analysis confirms the specific pattern of substitutions which indicate a series of tinkering mutations by trial and error method. For example, a series of substitution could be seen at the flexible regions of the α-helices where the Gly residues were substituted by N-and O-glycosylation sites (G446S, G496S of Omicron variant). Interestingly, the O-glycosylation sites are being substituted with N (S447N of Omicron variant and T501N substitution of SARS-CoV ? SARS-CoV-2), which were confirmed by LC-MS studies of SARS glycosylation profile (Shajahan, Archer-Hartmann, et al., 2020; Shajahan, Supekar, et al., 2020; Watanabe et al., 2020). Later variants evolved after the COVID-19 outbreak indicate the substitution of redundant N-glycosylation sites with the bulky Tyr (N501Y of Omicron variant) or Lys residues. Additionally, the Asp and Arg substitutions were observed more frequently. The Arg is known to interact with both aromatic and aliphatic side chains because of the unique gunanidium group which facilitates both the stacking and hydrogen bonding interactions. The polar negatively charged Asp substitutions may play role in stabilizing the complex by forming salt bridges and/or interact with the positively charged residues of ACE2. All the mutations discussed above are just one nucleotide distance away from the previous residue which simplifies our understanding of evolution. Moreover, the physio-chemical distance emphasize the necessity of the acquired substitution which was corroborated by the docking analysis. In the due course of evolution, the variants have shown increased binding affinity against the ACE2 receptor. Inspite of their ubiquitous and wide ranging role glycans are the least studied biomolecules, because of their complex structures which are quite heterogeneous and dynamic in nature making it a challenge to capture by the existing structural and biophysical techniques. This paper will serve as a precursor for the future in vitro and in vivo studies helping us to understand the dynamic aspects of the elusive glycans

## Abbreviations

ACE2: Angiotensin Converting Enzyme 2
CNI: Close-Neighbor-Interchange
CoV: Corona Virus
HADDOCK: High Ambiguity Driven Protein-Protein Docking
LC-MS: Liquid Chromatography - Mass Spectroscopy
Pangolin: Phylogenetic Assignment Of Named Global Outbreak Lineages
PGI: Protein-Glycan Interactions
RBD: Receptor Binding Domain
PPI: Protein-Protein Interactions
SARS: Severe Acute Respiratory Syndrome
SASA: Solvent Accessible Surface Area
SNP: Single Nucleotide Polymorphism
VOC: Variant Of Concern

## Acknowledgements

PKM acknowledges the National Institute of Technology Warangal for providing institute fellowship.

## Declaration

The authors declare no conflicts of interest.

## Author Contributions

The study was designed by TC, AM, SKG and PKM. The initial draft was prepared by PKM. The final manuscript was proofread by all the four authors.

---

**Figure S1:**
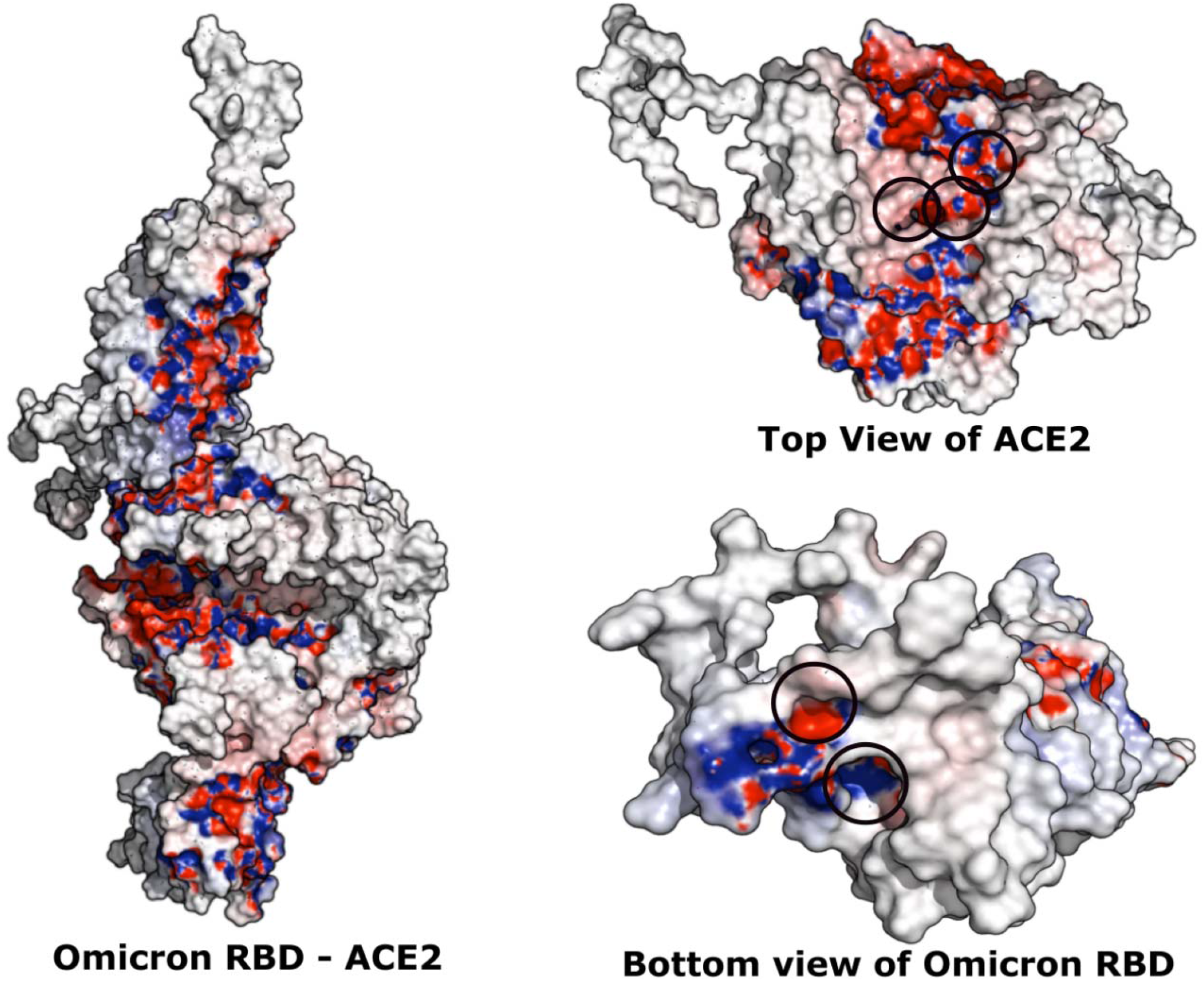
The electrostatic potential of the receptor binding domain of Omicron (Variant of Concern) with ACE2 receptor (left) of *Homo sapiens* are shown in surface representation. The top view of ACE2 receptor (right top) and bottom view of Omicron RBD (right bottom) are visualized for better understanding. The negative electrostatic potential are highlighted in red and the positive in blue.

